# Investigating orientation adaptation following naturalistic film viewing

**DOI:** 10.1101/2024.08.07.607115

**Authors:** Emily J A-Izzeddin, Reuben Rideaux, Jason B Mattingley, William J Harrison

## Abstract

Humans display marked changes to their perceptual experience of a stimulus following prolonged or repeated exposure to a preceding stimulus. A well-studied example of such perceptual adaptation is the tilt-aftereffect. Here, prolonged exposure to one orientation leads to a shift in the perception of subsequent orientations. Such a capacity to adapt suggests the visual system is dynamically tuned to our current visual environment. However, it remains unclear to what extent adaptation occurs in response to systematic features in naturalistic scenes. We therefore investigated orientation adaptation in response to natural viewing of filtered live-action film stimuli. Within a session, participants freely viewed 45 minutes of a film which had been filtered to include increased contrast energy within a specified orientation band (0°, 45°, 90°, or 135°; i.e., the adaptor). To measure adaptation effects, the film was intermittently interrupted to have participants perform a simple orientation judgement task. Having participants complete behavioural trials throughout the testing session, including 45 minutes of total adaptation time, allowed investigation of the accumulation of response biases and changes in such biases over the course of the session. We found participants exhibited stronger adaptation effects in response to cardinal adaptors compared to obliques. However, overall adaptation effects were weaker than those observed under typical tilt-aftereffect paradigms. Further, within a single session, adaptation effects developed inconsistently. The current findings therefore demonstrate a resistance to adaptation in response to naturalistic viewing conditions, suggesting barriers to understanding perceptual adaptation as experienced in nature.

## 1. Introduction

The key to any organism’s success is its ability to adapt to its environment. Humans, for example, are exposed to and interact with many different environments on a daily basis, behaviourally adapting to context-specific conditions with relative ease. This is no small feat, with the unique requirements of each environment ranging from complex factors that we are consciously aware of (e.g., understanding whether we need to change our clothing to better suit current weather conditions) all the way down to extremely basic perceptual factors that we have little to no awareness of (e.g., determining how well the distribution of basic visual features, such as orientation, match our prior expectations; A-Izzeddin et al., 2024; Frazor & Geisler, 2006; Torralba & Oliva, 2003; West et al., 2023). The extent to which each environment differs and our ability to cope with such variations suggests an extremely flexible and adaptive mechanism underlying our interactions with our environment.

At the level of visual perception, humans’ ability to adapt to specific visual features, such as orientation, has led to important insights into basic visuo-cognitive function. One of the most well-known approaches to understanding adaptation in this domain is the use of the classic “aftereffect” paradigm, initially described by Gibson and Radner (1937). Such paradigms are typically characterised by having participants fixate an oriented “adaptor” stimulus for several seconds. Subsequently, participants observe a “test” stimulus, positioned at a new orientation, and are asked to report the test stimulus’ orientation relative to a specified orientation (e.g., vertical). The hallmark finding in such paradigms is a perceptual “tilt-aftereffect”, occurring after the removal of the adaptor stimulus, whereby perception of a subsequent test stimulus’ orientation is altered relative to the adapted orientation (**Fig. 1A-B**; Calvert & Harris, 1988; Campbell & Maffei, 1971; Clifford et al., 2000; Gibson & Radner, 1937; Harris & Calvert, 1989; Mitchell & Muir, 1976; Morant & Harris, 1965). Specifically, if the test orientation is within ∼50° of the adaptor orientation, a repulsive effect is observed, whereby the test stimulus appears to be oriented away from the adaptor orientation (**Fig. 1B, positive values**; Clifford et al., 2000). Conversely, for test orientations greater than ∼50° relative to the adaptor orientation, an attractive effect is observed, whereby the test stimulus appears to be oriented towards the adaptor orientation (**Fig. 1B, negative values**; Clifford et al., 2000). The strength of such adaptation effects has been measured as a function of adaptation time, with results suggesting that the tilt-aftereffect builds up logarithmically in response to longer adaptation times (Greenlee & Magnussen, 1987; Magnussen & Greenlee, 1985; Magnussen & Johnsen, 1986), saturating after approximately an hour (Greenlee & Magnussen, 1987; Magnussen & Greenlee, 1985). Indeed, the fact that our perception is demonstrably impacted by an immediately preceding stimulus exemplifies our visual system’s capacity for highly specific short-term adaptation to its most recent input.

**Figure 1.**
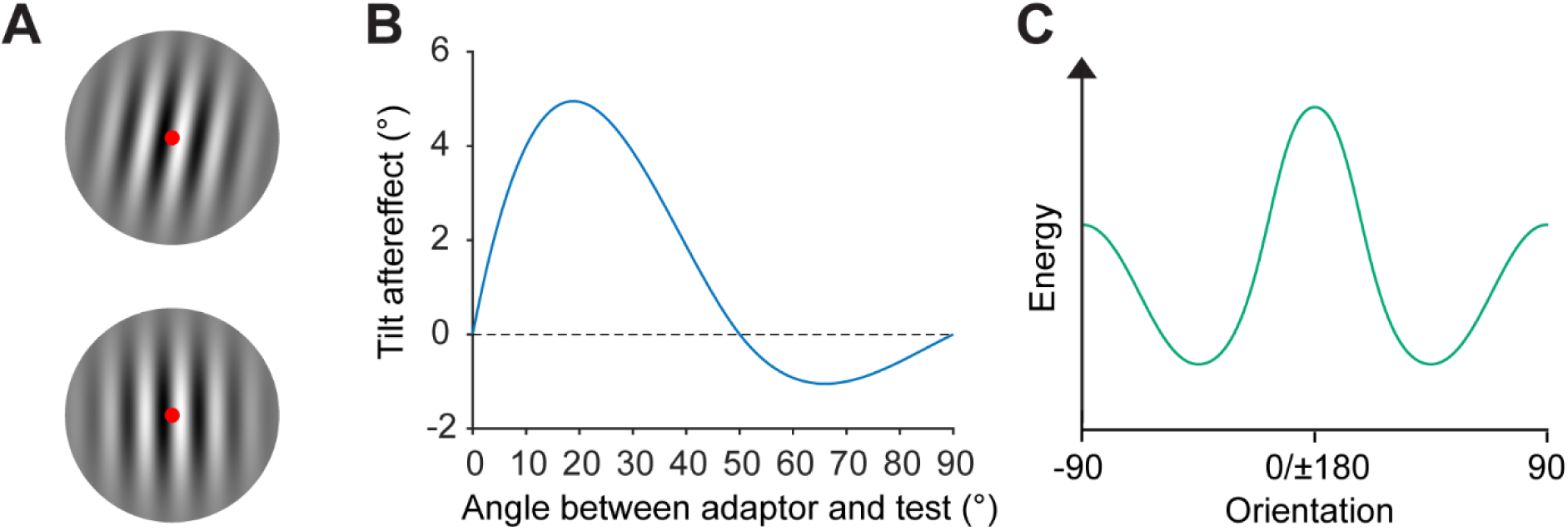
Tilt aftereffect and orientation prior overview. **A)** Example of the tilt aftereffect. Stare at the central red fixation dot of the top Gabor patch for 30 seconds, then shift your fixation to the fixation dot of the bottom Gabor patch. The bottom patch may appear to be tilted counterclockwise despite being vertical. This is the tilt aftereffect, which is a perceived repulsion away from the orientation of the adapting Gabor. **B)** Stereotypical aftereffects observed as a function of the difference in orientation between the adaptor and test stimulus. Positive values indicate a repulsive effect, and negative values indicate attraction (adapted from Clifford et al., 2000). **C)** Typical distribution of orientations observed in naturalistic images, with a dominance of cardinal orientations over obliques. This dominance has led to our prior for orientation information, resulting in the oblique effect.

Biologically, the tilt-aftereffect is thought to arise from the adaptation of neurons involved with processing the adaptor stimulus, which undergo response suppression after prolonged or repeated stimulation (Kohn, 2007). This response suppression leads to overall shifts in neural preferential tuning, resulting in a distorted representation of the subsequent test orientation relative to the adaptor (Kohn, 2007; Rideaux et al., 2023). The fact that such tuning shifts occur has been suggested to be of functional benefit to the observer, acting to dynamically tune the visual system to optimally process the visual information yielded from the current visual environment (Clifford et al., 2007; Kristjánsson, 2011).

Beyond short-term adaptation to immediate sensory input, we possess broader heuristics that we use to interpret the elements of our environment – expectations, or priors, thought to be built up over the course of our lives (A-Izzeddin et al., 2022; Series & Seitz, 2013; Summerfield & Egner, 2009) and even over evolutionary timescales (Geisler, 2008; Harrison et al., 2023). Of particular relevance to the tilt-aftereffect literature is our prior for orientation distributions. In nature, there is a dominance of cardinal orientations over oblique orientations (**Fig. 1C**; Coppola et al., 1998; Essock et al., 2003; Girshick et al., 2011; Hansen et al., 2003; Hansen & Essock, 2004; Keil & Cristóbal, 2000), a pattern reflected in observers’ greater sensitivity to cardinal orientations than obliques, otherwise known as the “oblique effect” (Appelle, 1972; Berkley et al., 1975; Campbell et al., 1966; Dakin, 2001; Dakin et al., 2009; Dakin & Watt, 1997; de Gardelle et al., 2010; Emsley, 1925; Girshick et al., 2011; Harrison et al., 2023; Pratte et al., 2016; Westheimer & Beard, 1998). This sensitivity has been observed under a multitude of experimental paradigms. However, the extent to which we can induce updates to this prior within the constraints of an experimental session has received relatively less attention in the literature, as opposed to typical short-term adaptation effects.

Importantly, the probing of long-term priors for orientations is facilitated by the use of complex stimuli. Complex stimuli allow for many cross-channel interactions, as in nature (Hansen & Hess, 2012; Webster & Miyahara, 1997). This poses a clear divergence from classic short-term adaptation investigations, which have typically involved a single adaptor and no conflicting information, likely isolating response channels in a manner that is very different to natural feature processing (Calvert & Harris, 1988; Campbell & Maffei, 1971; Harris & Calvert, 1989; Mitchell & Muir, 1976; but see Georgeson & Meese, 1997). Such constrained stimuli have therefore been inherently unable to investigate cross-channel interactions that could occur in response to more complex scenes. Further, there remains an open question regarding the timescales over which adaptation to complex stimuli might be elicited. For example, it is possible that participants’ initially demonstrate a repulsive adaptation pattern similar to that predicted by a typical tilt-aftereffect (Calvert & Harris, 1988; Campbell & Maffei, 1971; Clifford et al., 2000; Gibson & Radner, 1937; Harris & Calvert, 1989; Mitchell & Muir, 1976; Morant & Harris, 1965), but this might shift to reveal an attractive pattern, in line with biases consistent with our long-term orientation priors (Harrison et al., 2023).

Given the relative lack of investigations of adaptation using complex stimuli, it remains unclear if naturalistic distributions of ordered spatial structure that invoke high-level object and scene representations can elicit low-level adaptation effects. Naturalistic image stimuli pose an ideal candidate for probing adaptation in response to naturalistic distributions of features. Naturalistic images convey complex distributions of features, addressing the key concern surrounding typical approaches taken to understanding visual adaptation. Additionally, naturalistic images serve as a useful proxy for the real world, representing the features we observe in nature day to day, for which we have accumulated long-term priors (Geisler, 2008; Harrison, 2022; Simoncelli, 2003; Simoncelli & Olshausen, 2001; Theunissen et al., 2001). Relevant to the existing adaptation literature, naturalistic images convey the same distribution of orientations (**Fig. 1C**) and spatial frequencies that the human visual system is tuned to (David et al., 2004; Harrison, 2022; Olshausen & Field, 1996; Simoncelli & Olshausen, 2001). As such, naturalistic images present an ideal candidate for assessing observers’ capacity for naturalistic adaptation.

Beyond the use of naturalistic images, some studies have gone so far as to employ live-action films to better simulate naturalistic viewing conditions. For example, Dorr and Bex (2013) investigated the interplay between visual sensitivity and eye movements, having participants perform a detection task for a contrast increment target embedded in live-action films. Similarly, Wallis et al. (2015) investigated luminance contrast sensitivity using a target detection paradigm, similarly embedded in a live-action film. Of most relevance, Bex et al. (2009) investigated participants’ contrast sensitivity function after undergoing adaptation to live-action film stimuli. Bex et al. found that, compared with when no adaptor stimulus was presented, contrast sensitivity was reduced for low spatial frequencies following live-action film adaptation. As such, Bex et al. demonstrate the capacity to elicit adaptation effects for contrast sensitivity under natural viewing conditions. However, the aforementioned investigations have focused on contrast sensitivity in conjunction with live-action film stimuli, leaving open the question of whether adapted orientation sensitivity can be elicited from live-action film.

Therefore, in the current study, we investigated orientation adaptation by filtering live-action film stimuli. We filtered a film to have uniform orientation energy across all spatial frequencies, except for at relatively low spatial frequencies, which were filtered to contain only one specified orientation (i.e., the adaptor orientation; **Fig. 2**). Specifically, participants were shown the film in halves across separate days. Each film half (45 minutes of footage) was subjected to one of four filtering conditions, such that each participant saw 45 minutes of the film with only one cardinal orientation (i.e., 0° or 90°) present at relatively low spatial frequencies, and the other 45 minutes with only one oblique orientation (i.e., 45° or 135°). Differences between cardinal and oblique adaptors were of interest because the anisotropic encoding of orientations could yield differential adaptation effects (Campbell et al., 1966; Furmanski & Engel, 2000; Li et al., 2003; Maloney & Clifford, 2015). The filtered film was intermittently interrupted to have participants perform a basic orientation judgement task, indicating whether a centrally presented Gabor was rotated clockwise or counterclockwise relative to a peripheral bar. By having participants freely view a film stimulus, we presented naturalistic images in an engaging manner that more closely simulates participants’ real-world experience than typical tilt-aftereffect paradigms. Additionally, having participants view the film for 45 minutes over the course of a session allowed us to investigate the timeline of any adaptation that occurred. Therefore, the use of such complex stimuli in conjunction with a clockwise/counterclockwise orientation judgement task allowed us to measure participants’ degree of adaptation to orientations presented under more naturalistic viewing conditions over time.

**Figure 2.**
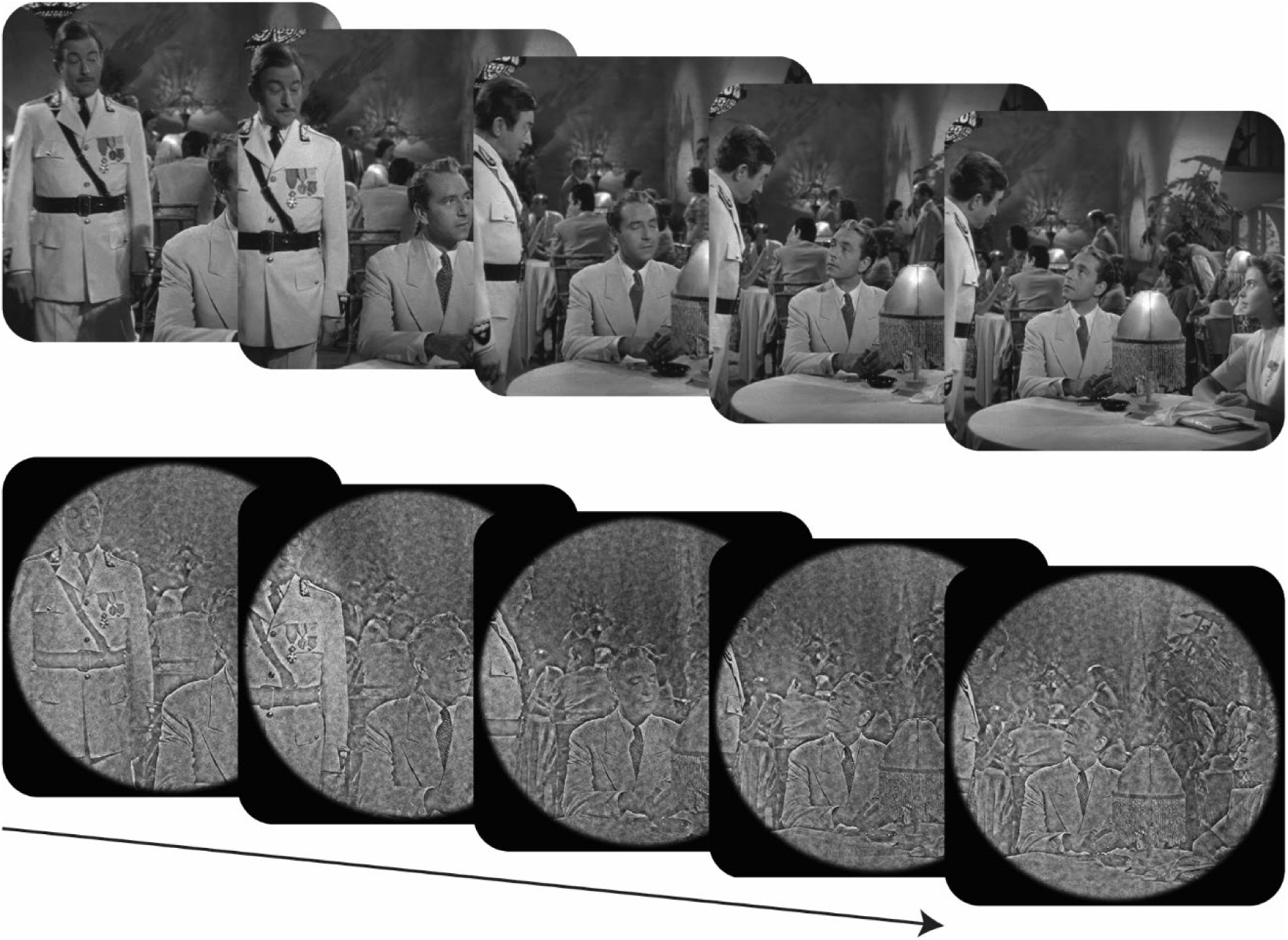
Adaptation in response to naturalistic image stimuli. In the current study, we presented participants with filtered film stimuli – in this case, from the film *Casablanca*. The top row are original film frames. Stimuli used in the experiment (bottom row) were filtered to only have one orientation present at relatively low spatial frequencies (1-4cpd) – 0° in this case. Using this stimulus, we assessed participants orientation adaptation in response to naturalistic images. Note: spatial frequency filters were based on actual stimulus presentation size. Hence, examples here will be distorted by reducing the size of the images.

## 2. Methods

### 2.1. Participants

Twenty-eight participants took part in the current study. Four participants were excluded – three failed the baseline data quality check (see **Section 2.4.3. Practice and baseline testing**), and one participant withdrew. Hence, 24 participants’ datasets were included in analyses. Participants had varying degrees of experience participating in psychophysical experiments, and all participants were naïve to the purpose of the experiments. All participants provided informed consent to participate in the current study. Ethics approval was granted by The University of Queensland (Medicine), Low & Negligible Risk Ethics Sub-Committee.

### 2.2. Stimuli

#### 2.2.1. Video stimuli

*Casablanca* was used as the source film for stimulus generation (Curtiz, 1942; the film is dedicated to the public domain under CC0). We extracted each frame from the digitised film as individual image files. Each frame was first whitened such that there was equal energy at all orientations and spatial frequencies. Subsequently, a 1/f spectrum was applied to the frames. We then applied a combined orientation and spatial frequency filter to the frames. The spatial frequency filter was flat-topped, uniformly covering 1-4cpd, with cosine edges falling to zero over half an octave. The orientation filters were raised cosine filters with a period of 45°. Four versions of the orientation filter were generated, such that four versions of the film were produced that had each frame filtered to contain only 0°, 45°, 90°, or 135° orientations (i.e., the possible adaptor orientations) at the specified spatial frequencies (**Fig. 3A**). Filtering at 1-4cpd meant that filtered orientations were in a frequency range that approximately corresponds to the peak tuning of humans’ contrast sensitivity function (Bex et al., 2009). Finally, each final frame’s total energy was matched to that of the original unfiltered frame, thereby retaining the overall energy dynamics of the original film. We then cropped frames to a 14.30°-diameter circle – the maximum possible diameter given the film frame size, display resolution, and viewing distance. We cropped frames using a circular aperture to remove cardinal orientation cues that would have otherwise been introduced at the edges of the stimulus.

**Figure 3.**
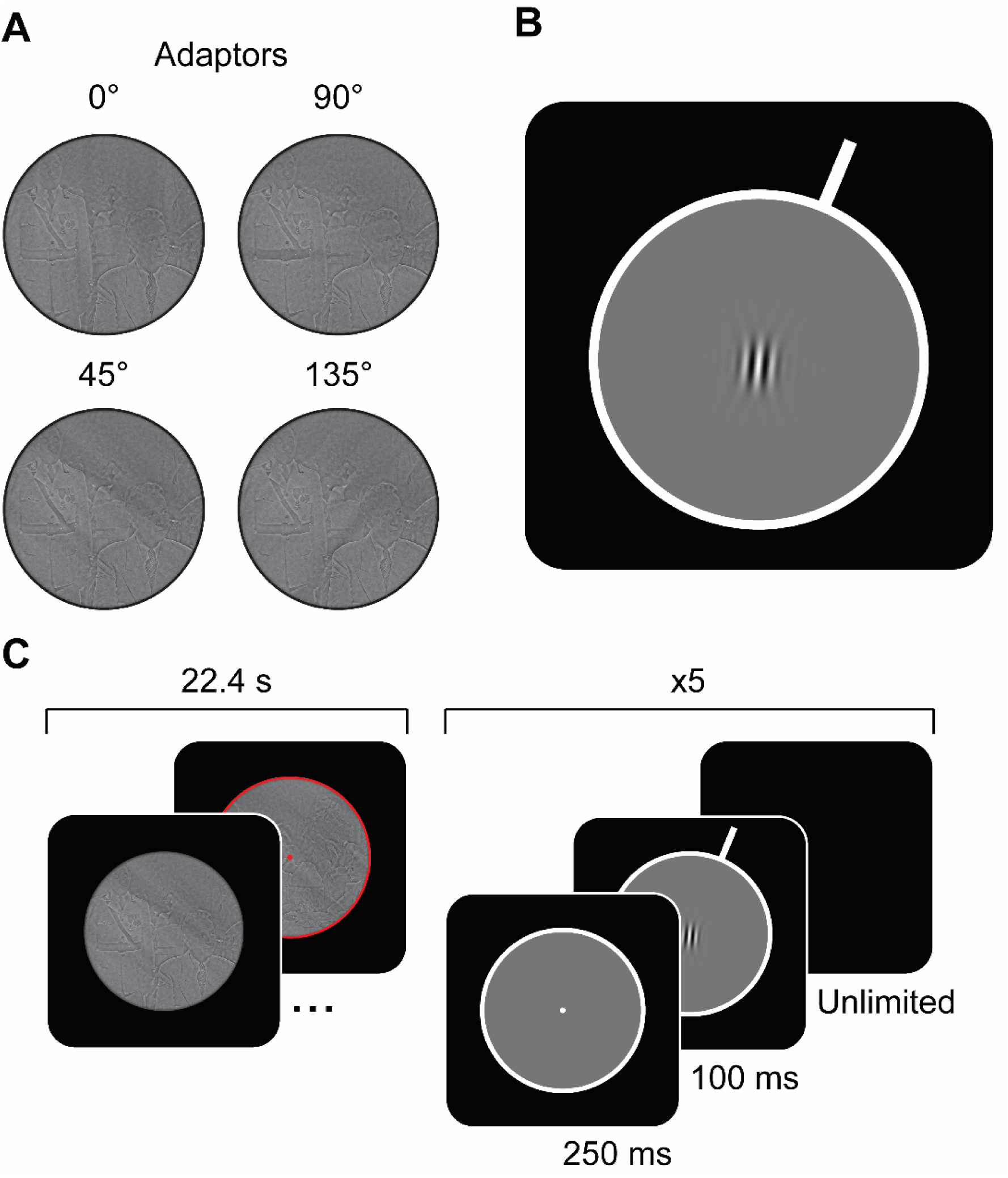
Overview of stimuli. **A)** An example of the same frame from the film, *Casablanca*, after passing through one of four orientation/spatial frequency filters, resulting in all orientations at 1-4cpd being removed, except for 0°, 90°, 45°, or 135°. Note: Images displayed here are smaller than those displayed to participants, hence filtering visible here will not correspond to 1-4 cpd. **B)** Example of the perceptual task stimulus. Observers reported whether the peripheral white bar (the standard stimulus) was oriented clockwise or anti-clockwise relative to the central Gabor patch (the test stimulus). The white bar was positioned peripherally, outside of the area taken up by the adaptor stimulus. Because participants were viewing the video itself, the standard should not be subject to the same level of adaptation as the central Gabor (which did occupy the same space as the preceding adaptor). **C)** Participants viewed a 22.40 s filtered film clip, followed by five trials wherein participants were asked to indicate whether the central grating is tilted clockwise or counterclockwise relative to the standard stimulus (i.e., the protruding white bar). This structure repeated 120 times in each session until participants had watched 44.80 minutes of the film, *Casablanca*, and had completed 600 trials.

After filtering, film frame images were re-combined to make a series of 22.40s video clips, generating a total of 240 clips that amounted to 89.6 minutes. We chose the current clip length to break up the film into enough segments to allow for fewer behavioural trials (i.e., five) between each clip, and to allow for an even number of clips in each session (i.e., 120; see **Section 2.4.4., Adaptation**). We also included the associated audio files in generated clips to allow participants to listen to the film while watching.

At the end of a clip, participants immediately completed behavioural task trials. As such, we included a cue to warn participants that the clip was about to end, and a trial was about to start. The cues were a 0.12°-stroke red border that appeared around the edge of the stimulus, as well as a central 0.11° red fixation point. The cues were presented for the final second of each clip (**Fig. 3C**, left).

#### 2.2.2. Perceptual task stimuli

Participants watched the filtered video clips in sequence, with a break between each clip to collect behavioural responses. All stimuli were presented on a black background, with a centrally positioned 14.30° grey circle, framed by a white border (stroke = 1.00°; **Fig. 3B**). The fixation point, when shown, was presented centrally, subtending 0.11°. The trial stimulus consisted of two components: the standard stimulus and the test stimulus. The standard stimulus was made up of one white bar, 2.24° in length and 1.00° stroke, extending from the white border surrounding the central grey circle at one of four orientations (-67.5°, -22.5°, 22.5°, or 67.5°). The test stimulus was defined in the frequency domain by a two-dimensional raised cosine function, with the radial axis corresponding to orientation and the tangential axis corresponding to spatial frequency. The full-width half weight of the filter was 45° for orientation, and 1.148 cpd for spatial frequency. Note that the resulting test stimulus’ spectrum was therefore well-equated with the profile of the adapting stimuli. The test stimulus was drawn centrally. Gabors were presented at 100% RMS contrast (with recent papers demonstrating tilt aftereffects with high-contrast test stimuli; Gurbuz & Boyaci, 2023; Knapen et al., 2010; Nakashima & Sugita, 2014). The grating had a spatial frequency of 2 cpd and a 0° phase. Gabors on each trial were drawn at one of seven orientation offsets relative to the standard stimulus (-10°, -5°, -2.5°, 0°, 2.5°, 5°, or 10°).

### 2.3. Apparatus

Stimuli were generated on a ThinkStation P520 computer (running Windows 10 Enterprise) with the Psychophysics Toolbox (3.0.17; Brainard, 1997; Pelli, 1997) for MATLAB (R2018a). Stimuli were presented on a 24-inch Asus VG428QE 3D monitor with 1920 x 1080-pixel resolution and a refresh rate of 100 Hz. A gamma correction was applied to the display, assuming a gamma of 2. As noted by Bex et al. (2005), small errors in gamma are inconsequential to the representativeness of the naturalistic images. A cardboard circular mask was superimposed on the monitor to block out orientation cues conveyed by the edges of the screen. Film audio was provided through Sony WH-CH710N Wireless Noise Cancelling Headphones (connected via AUX cable) at participants’ desired volume. Testing sessions were completed in a testing booth with the lights off.

### 2.4. Task design

The following sections will outline the different elements of the experiment design. For a flow chart summarising the key session, trial, and task elements of the experiment, please see **Supplement S.1**.

#### 2.4.1. Condition counterbalancing

Each participant watched one half of the film with a cardinal adaptor (0° or 90°) and the other half of the film with an oblique adaptor (45° or 135°; 45 minutes per half). Combinations and order of adaptors were counterbalanced across participants. Participants were shown only one cardinal and one oblique adaptor to maximise the adaptation time and trials for each condition. Counterbalancing condition assignment allowed us to investigate adaptation to all adaptor conditions.

We wanted to ensure that participants saw standard stimuli at orientations that were equidistant from the two adaptors observed across sessions. Such equidistance is important as the strength of the tilt-aftereffect is known to be highly dependent on the orientation of the test stimulus relative to the adaptor orientation (**Fig. 1B**; Clifford et al., 2000). Further, selecting standard orientations to be equidistant from adaptors meant that only the orientation of the adaptor changed across sessions.

#### 2.4.2. General trial structure

Participants completed five perceptual task trials between film segments. Participants were alerted to the impending start of a perceptual trial via the appearance of a red border and fixation point for the last second of the film clip (see **Section 2.2.1**., **Video stimuli**). On each behavioural trial, participants indicated whether the central test stimulus was rotated clockwise or counterclockwise relative to the standard stimulus (**Fig. 3C, right**). Trials began with a central white fixation point, presented for 250 ms, after which the stimulus was shown, with the standard stimulus at its given orientation for that trial and the test stimulus at its given orientation offset relative to the standard. The standard/test stimulus was presented for 100 ms, followed by a blank screen which remained until a response was recorded. Participants were instructed to use the left or right arrow keys to make their response. During trials, participants were instructed to fixate on the test stimulus when presented. Critically, the standard stimulus was positioned to be outside of the space occupied by the film stimulus (i.e., the adapting region) to minimise the impact of adaptation on perception of the standard.

Participants were not subject to eye tracking and were free to move their eyes while watching the video stimuli. As such, the retinotopically-mapped adaptation region likely extended beyond the confines of the spatially-mapped film stimulus area. Nonetheless, on average, we expect participants to look towards the centre of the film stimulus area (Dorr et al., 2010). Therefore, the level of adaptation experienced in the retinotopic areas that correspond to the standard stimulus should be weaker than that experienced foveally. Further, the brevity of the test and standard stimulus presentation (100 ms) would pose difficulty in attempting to foveate both stimuli. Hence, such a tactic would likely lead to decrements in task performance. Given participants’ performance is assessed after the initial testing session (see **Section 2.4.3., Practice and baseline testing**), it is unlikely such saccades can explain performance observed.

#### 2.4.3. Practice and baseline testing (Session 1)

Prior to adaptation, in a separate initial session, participants completed 588 practice trials without viewing any film stimuli. For the first 84 trials, test stimulus offsets relative to the two standard orientations used were doubled (i.e., to -20°, -10°, -5°, 0°, 5°, 10° and 20°). The final 504 practice trials had offsets matching the general trial structure (i.e., -10°, -5°, -2.5°, 0°, 2.5°, 5°, or 10°). Participants received feedback in the form of a red or green (incorrect and correct, respectively) fixation point, presented centrally for 750 ms following response. Following feedback, the next trial would begin.

Practice trials were followed by 600 experimental trials to quantify participants’ baseline orientation biases as measured by the task. No feedback was given for baseline trials. Baseline trials had test stimulus offsets matching the general trial structure described above. Given the limits placed on trial numbers due to time constraints, and that smaller offsets were expected to yield noisier responses, we had participants complete more trials for smaller offsets than larger offsets in order to constrain the psychometric fits described below. Specifically, for each of the two standard stimulus orientations, there were 20 trials each for -10° and 10° offsets, 40 trials each for -5° and 5° offsets, and 60 trials each for -2.5°, 0°, and 2.5° offsets.

After a participant completed the baseline trials, we conducted a data quality check to evaluate if their performance was above chance. The participant’s baseline data were fit with two generalised linear regression models using MATLAB’s “fitglm” function. The first model was an intercept-only model, and the second included intercept and slope parameters, using a logit link function (i.e., a psychometric function). The log likelihoods for each model were compared and participants “passed” the data quality check if the psychometric function was a significantly better fit (*p* < 0.05) than the intercept-only model. Specifically, we computed the likelihood ratio chi-square statistic by first doubling the difference in log likelihoods between the two models. We then calculated the chi-square cumulative distribution function at this difference value, subtracted from 1, and interpreted the resulting value as a *p*-value (Quandt, 1958; Taboga, 2021).

#### 2.4.4. Adaptation (Sessions 2-3)

Participants’ second and third sessions each started with a block of 210 practice trials. Practice trials had test stimulus offsets matching the general trial structure. For the first 14 trials (one trial per standard stimulus orientation and test stimulus offset combination), participants received feedback. For the last 196 trials, no feedback was given, and there was an even distribution of trials across standard stimulus orientation and test stimulus offset combinations (i.e., 14 trials per combination). Responses for the last 196 trials were included in participants’ baseline dataset for analyses.

Practice trials were followed by the adaptation component of the session. During each adaptation session, participants observed 120 filtered video clips (each 22.40s in length) taken from the film, *Casablanca,* intermingled with trials. Clips were shown in the order they are presented during the film, such that participants were able to follow the original narrative structure. Participants completed 600 trials in each adaptation session. Trials followed the general trial structure and were split into 120 blocks of five, with each block preceded by a video clip (**Fig. 3C**). In the first adaptation session, participants watched the first 44.80 min of *Casablanca* with either an oblique adaptor or cardinal adaptor applied. In the second session, participants viewed the second 44.80 min with the opposite adaptor-type to that used in the previous session (counterbalanced across participants). Participants viewed the video with audio played through headphones.

At the end of each block of five trials, participants were shown a countdown timer, which was a circle that gradually filled like a timer, and gave participants an opportunity to have a break if needed. If participants requested a break, the screen went black, and participants could re-start when ready. Participants were asked to only take breaks when necessary. No participants left the dark testing booth during their break, minimising their exposure to the statistics of the broader laboratory during their breaks.

At the end of the final session, participants had viewed 89.60 min of *Casablanca*. However, there were an additional ∼10 mins of the film that was not filtered. Participants were given the option to view this after completing the experiment, with the majority choosing to do so, acting as a potential indicator that participants were invested in attending to the film. We note that in this time, unprompted, several participants indicated that they did *not* realise that the film they had watched during the experiment was altered until seeing the unaltered final 10 mins of the film. These participants commonly suggested they had attributed the altered film’s appearance to its age.

### 2.5. Analyses

#### 2.5.1. Generalised linear multilevel modelling

Participants’ performance was modelled using a generalised linear multilevel model (GLMM) framework. The GLMM was fit to all of the data using MATLAB’s fitglme() function. Participant responses were coded as 0 and 1, indicating that the test stimulus was perceived to be oriented to the left or right of the standard, respectively. The model included four predictors based on trial-by-trial conditions. Specifically, the first predictor was the offset of the test stimulus relative to the standard (i.e., -10°, -5°, -2.5°, 0°, 2.5°, 5°, or 10°). The second predictor was the adaptor condition (i.e., 0°, 45°, 90°, or 135°). The third predictor was the standard orientation (i.e., -67.5°, -22.5°, 22.5°, or 67.5°). The fourth predictor was the interaction between adaptor and standard orientation conditions – implemented because the combination of standard orientations a given participant saw was dependent on the combination of adaptor conditions they observed across sessions. For the baseline condition, where no adaptor was used, an additional condition label of ‘-999’ was added to the adaptor condition predictor. All predictors were categorical. The probability of a right response was predicted as:

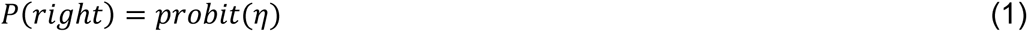

where *probit* represents the probit link-function, and:

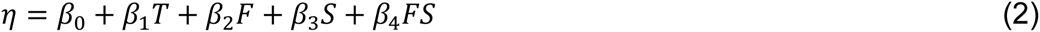

where β_0_ is the intercept term, β_1_ is the weight of the test stimulus offset relative to the standard, *T*, β_2_ is the weight of the adaptor condition, *F*, β_3_ is the weight of the standard orientation, *S*, and β_4_ is the weight of the adaptor/standard orientation interaction, *FS*. To partially pool coefficient estimates across participants, the GLMM included participant as a random effect. We describe this approach in more detail in Rideaux et al. (2022). For visualisation of fits (e.g., **Fig. 4**), we found the GLMM estimate for each adaptor by standard orientation condition combination, representing the conditional intercept, which we interpret as observers’ response bias for a given condition.

**Figure 4.**
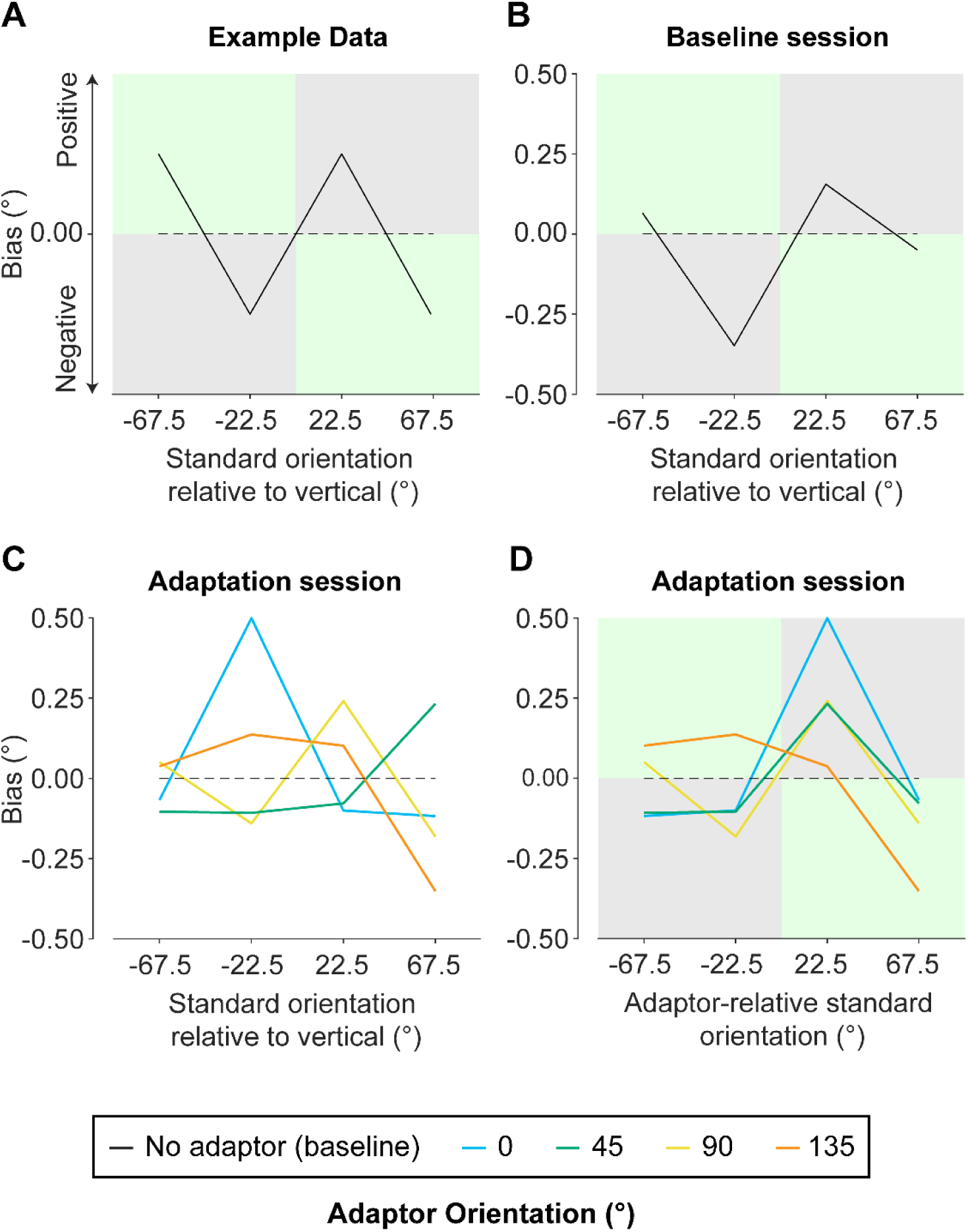
Bias data from baseline and adaptation sessions. **A)** Example data demonstrating what pattern of results should be found with a typical cardinal-repulsion effect. The standard orientation (relative to vertical) is on the x-axis and bias shift on the y-axis. Grey panels indicate repulsive response zones and green panels indicate attractive response zones (relative to vertical). The black dotted line is included to highlight deviations from zero bias. **B)** GLMM fits of participants’ baseline bias data with the standard orientation (relative to vertical) on the x-axis and bias shift on the y-axis. **C)** GLMM fits of participants’ adaptation bias data relative to baseline for each adaptor orientation (see bottom legend). Absolute standard orientation (relative to vertical) is on the x-axis and bias shift on the y-axis. Here, panels indicating attractive vs repulsive biases are omitted as whether a bias is attractive or repulsive will depend on the specific adaptor orientation and standard orientation combination. Note, these data have had baseline data subtracted to view biases as a result of adaptation alone. **D)** Here, the same data from Panel B are re-plotted, with standard orientations re-coded to be relative to the adaptor orientation. This normalisation allows consistent interpretation of whether responses are attractive/repulsive for a given standard orientation. As such, grey and green panels are used again to indicate repulsive and attractive biases, respectively. Note: Biases shown are derived from combining multiple parameters, preventing plotting of their confidence intervals directly (see **Supplement S.2** for full model output).

#### 2.5.2. Exploratory sequential analyses

Because we were interested in how adaptation effects emerge and evolve over prolonged viewing of systematically skewed image statistics, we conducted an exploratory sequential analysis to investigate changes in response bias across trials during testing sessions. For example, it is possible that participants’ initially demonstrate a repulsive adaptation pattern similar to that predicted by a typical tilt-aftereffect (Calvert & Harris, 1988; Campbell & Maffei, 1971; Clifford et al., 2000; Gibson & Radner, 1937; Harris & Calvert, 1989; Mitchell & Muir, 1976; Morant & Harris, 1965), but this might shift to reveal an attractive pattern, in line with biases consistent with our long-term orientation priors (Harrison et al., 2023). Sequential analyses were conducted for each standard orientation tested. The possible standard orientations a participant saw were one of -22.5° and 22.5° and one of -67.5° and 67.5°. When participants make a left/right response relating to whether the test is rotated to the left/right of the standard, the response will be attractive or repulsive depending on the standard orientation. For example, if we consider whether a response is attractive relative to the nearest cardinal, a leftward bias for the -22.5° standard orientation condition is attractive towards the nearest cardinal (i.e., vertical). However, a leftward bias for the 22.5° condition would be repulsive. However, this mapping of attractive vs repulsive responses arising from adaptation will change depending on the orientation of the adaptor. Therefore, to allow for consistent interpretation, data were normalised such that standard orientations were re-coded to indicate their orientations relative to the adaptor orientation.

Response bias is equivalent to the average response made across trials. Hence, to measure changes in response bias for a single relative standard orientation, we calculated the cumulative average response with each new trial. For example, if participants responded with “right” on the first trial (coded as 1 for this analysis), then the response bias for the first trial will perfectly equal 1. Then, for the second trial, we take the average of this second trial as well as the first trial. Should participants respond “left” for the second trial (coded as -1 for this analysis), then the response bias will average out to 0. This process continues until the last trial in a given session/for a particular standard orientation, which calculates the average response across all trials (300 in total per session/standard orientation). Because there were equal numbers of trials where the target was offset to the left and right of the standard, deviations from 0 indicate an overall response bias. Put differently, should participants display no response bias across the session, the sequential analysis should quickly even out to roughly 0 and remain consistent across the sequence.

We were interested in measuring if participants’ response bias at the start of the session differs to their response bias at the end of the session. To investigate this, we ran the same sequential analyses as described above with trials in reverse order. In this case, we start with the last trial, working backwards to the first trial, which calculates the average response across all trials. Should participants display a different response bias at the start of the session as compared with the end, then the cumulative response biases should differ between the forward and reverse accumulations.

For both the forward and reverse accumulations, we generate a bias accumulation trace that plots the cumulative bias across trials (**Fig. 5; pink and green lines**). To formally quantify whether there was a significant difference between forward and reverse cumulative biases, we conducted permutation analyses to generate a null distribution with associated confidence intervals for each relative standard orientation condition (Efron & Tibshirani, 1993). Specifically, 1000 permutations of the sequential analysis were conducted. Each permutation involved randomising whether a given trial’s cumulative response bias was calculated in the forward or reverse direction. For example, if the 10^th^ trial was randomised to be forward-coded, then the cumulative response bias would be based on the first 10 trials participants completed. Alternatively, if the 10^th^ trial was randomised to be reverse-coded, then the cumulative response bias would be based on the last 10 trials participants completed. Permutation data for a given relative standard orientation condition were then used to calculate 95% confidence intervals by finding the .025^th^ and .975^th^ quantile of the permutation data for each trial (**Fig. 5; black lines**).

**Figure 5.**
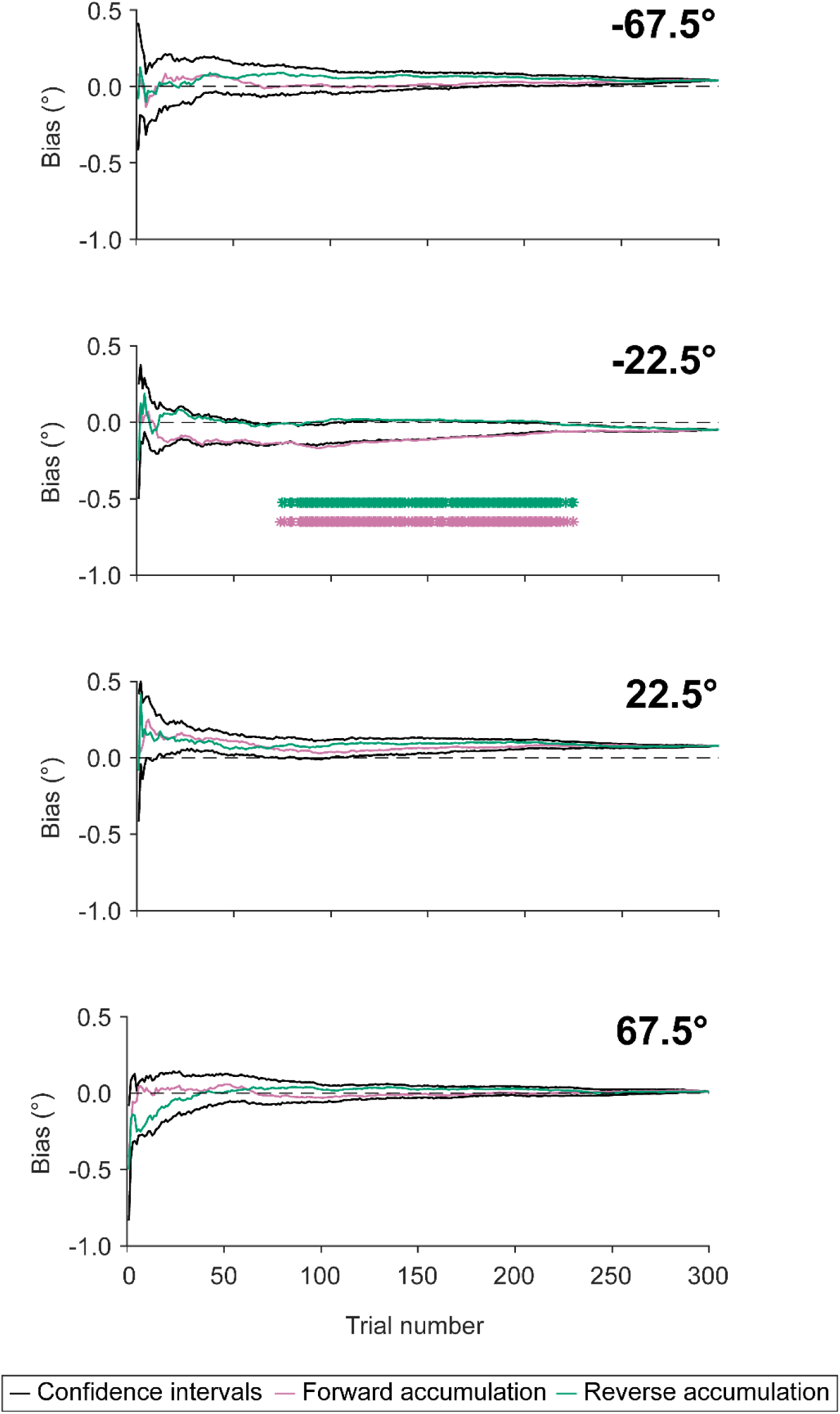
Mean bias accumulation data for adaptation sessions at each individual standard orientation (relative to the adaptor orientation), collapsing across adaptor conditions. Accumulated bias is calculated by taking the cumulative mean response spanning from trial 1 to 300. This was done in forward (i.e., ascending from 1; pink) and reverse (i.e., descending from 300; green) directions, however both are plotted in the same timeline to visualise differences in biases at the start vs end of the adaptation session. Black lines represent 95% confidence intervals, calculated from null distributions obtained from permutation analyses. Asterisks represent instances where the mean bias accumulation falls outside of the confidence intervals, indicating statistically significant differences between forward vs reverse accumulations at that particular time point.

## 3. Results

We quantified the extent to which prolonged adaptation to orientation-filtered videos influences subsequent orientation judgements. Participants completed three testing sessions in which we measured their baseline orientation bias, and then following adaptation to filtered video clips. The effects of adaptation were measured using a behavioural task in which participants judged whether a centrally presented Gabor patch was tilted clockwise or counterclockwise relative to a peripherally presented oriented bar. The first session measured participants’ “baseline” orientation bias prior to viewing the adaptor stimulus. The second and third sessions were similar in design to a typical tilt-aftereffect experiment, where participants were shown clips of a film which had had its orientation information altered (i.e., the adaptor/top-up stimuli), which were separated by Gabor judgement trials. Specifically, each participant completed a session with a cardinal (0° or 90°) adaptor, where the film had been filtered to have only one specified cardinal orientation present at relatively low spatial frequencies, and another session with an oblique adaptor (45° or 135°). The inclusion of a baseline session allowed us to assess each individual’s relative degree of adaptation to the adaptor stimuli. We assessed adaptation using a method of constant stimuli approach, calculating shifts in bias of participants’ clockwise/counterclockwise judgements relative to the orientation of the adaptor.

### 3.1. Adaptation effects in response to naturalistic stimuli are minimal

We fit a GLMM to our response data using the offset between the gabor and standard orientation (i.e., ±10°, ±5°, ±2.5°, and 0°) as well as the interaction between adaptor orientation (i.e., 0°, 45°, 90°, and 135°) and standard orientation (i.e., ±22.5° and ±67.5°) as predictors, with participants included as a random effect. The implementation of this GLMM is described in depth in **Section 2.5.1., Generalised linear multilevel modelling**. Here, we first outline the qualitative pattern of response biases observed for the baseline condition, followed by the bias pattern observed following adaptation. Finally, we outline the statistical output of GLMM fits.

We found repulsion away from the nearest cardinal axis in the baseline session, in line with previous oblique effect findings (Appelle, 1972; Taylor & Bays, 2018; **Fig. 4A-B**). Specifically, if we consider response biases relative to vertical, we found a repulsive bias for standard orientations close to vertical (i.e., ±22.5°). We found an attractive bias for standard orientations that deviate from vertical by ±67.5. An attractive bias in this instance is expected, given that ±67.5° is closer to horizontal than vertical. Hence, in the case of the ±67.5° standard orientation conditions, an attractive response bias relative to vertical is indicative of an overall repulsion away from the nearest cardinal orientation, horizontal.

We subtracted the baseline bias from each corresponding adaptor condition to assess the impact of adaptation alone on response bias. We found a relatively consistent shift in this bias pattern across adaptor orientations (**Fig. 4C-D**). Specifically, we observed repulsion for the closest relative standard orientations (i.e., ±22.5°) across all adaptor conditions. One exception to this effect is noted for the 135° adaptor condition, seen at the relative standard orientation of -22.5° (**Fig. 4D, orange**). Further, we found attraction for one of our largest relative standard orientations (i.e., 67.5°). However, for the -67.5° standard orientation, there was a less consistent attraction effect, which was only found for the 0° (**Fig. 4D, blue**) and 90° (**Fig. 4D, yellow**) adaptor orientations.

The GLMM revealed two predictor combinations were significant (see **Supplement S.2** for full model output). The first was a repulsive effect for the 0° adaptor orientation at the relative standard orientation of -22.5°. The second was an attractive effect for the 90° adaptor orientation at the relative standard orientation of 67.5°. These significant predictor combinations suggest stronger adaptation effects for cardinal-only adaptors, with weaker effects observed for oblique adaptors.

Overall, the strength of observed adaptation effects are much smaller than what we expected based on a classic tilt-aftereffect paradigm (Clifford et al., 2000). It is possible that the embedding of adaptor stimuli in more naturalistic surrounds acts to reduce adaptation effects or require longer exposure times to elicit strong adaptation effects. Alternatively, there could have been changes in adaptation throughout the testing sessions, mitigating the appearance of bias averaged across all trials, which we investigate below.

### 3.2. Adaptation does not reliably change within a testing session

Beyond investigating the difference in response bias between baseline and the end of adaptation, we were further interested in how this response bias changes over time as participants undergo adaptation. For example, it is possible that participants’ initially demonstrate a repulsive adaptation pattern similar to that predicted by a tilt-aftereffect (Calvert & Harris, 1988; Campbell & Maffei, 1971; Clifford et al., 2000; Gibson & Radner, 1937; Harris & Calvert, 1989; Mitchell & Muir, 1976; Morant & Harris, 1965), but this might shift to reveal an attractive pattern, in line with biases consistent with our long-term orientation priors (Harrison et al., 2023). We therefore investigated whether adaptation effects change over the course of a given session. In this section, we first outline the bias patterns observed in participants’ data. Following, we discuss the inferential approach taken to interpret the analysis’ results.

To investigate the time course of observed adaptation effects, we analysed the accumulation of participants’ response biases across time. The implementation of this analysis is described in depth in **Section 2.5.2., Exploratory sequential analyses**. In brief, this was done by taking the cumulative average response from trial to trial to see how participants bias shifts over time (**Fig. 5**). To assess whether there is a difference in adaptation effects at the start vs end of adaptation, accumulation analyses were run with responses in forward (**Fig. 5, pink lines**) and reverse order (**Fig. 5, green lines**). Separate analyses were conducted for each relative standard orientation to allow for consistent interpretation of whether a calculated bias is attractive or repulsive relative to the adaptor orientation. Comparing standard orientation conditions, there was noted variability in the time course of forward vs reverse bias accumulation. For example, the 22.5° relative standard orientation condition displays greater consistency in forward vs reverse traces than the -22.5°.

We then tested for statistically significant deviations between forward vs reverse accumulations in each relative standard orientation condition. To do this, we conducted permutation analyses on the time course data to generate a null distribution with associated 95% confidence intervals. The generation of confidence intervals allowed visualisation of when participants’ accumulated biases in forward and reverse directions are significantly different from one another (**Fig. 5, black lines**). We found significant deviation of response bias in the forward vs reverse direction only when the relative standard orientation is -22.5° (**Fig. 5, second plot from the top**). The lack of consistency in bias accumulation effects further suggests that embedding adaptor stimuli in more naturalistic surrounds may act to reduce adaptation effects and/or requires longer exposure times to elicit.

## 4. Discussion

We investigated visual adaptation to altered orientation statistics embedded in naturalistic stimuli. Participants watched films filtered to contain adaptors at a specific orientation and intermittently completed an orientation judgement task to measure shifts in response bias. We found stronger adaptation effects for cardinal adaptor orientations, with weaker effects observed for oblique adaptor conditions, and weak effects overall as compared with typical tilt-aftereffect studies. Further, we found adaptation effects do not consistently accumulate or fluctuate within a single testing session. Nonetheless, results of the current study suggest that adaptation, while limited, is possible to elicit in response to complex naturalistic stimuli to some extent. The small adaptation effects we did find are qualitatively similar to adaptation effects in response to isolated orientated contrast (Calvert & Harris, 1988; Campbell & Maffei, 1971; Harris & Calvert, 1989; Mitchell & Muir, 1976). Additionally, evidence has been found for orientation adaptation aftereffects in response to windowed naturalistic image regions with strong orientation cues (Dekel & Sagi, 2015). However, supporting evidence from the current study should be interpreted with caution due to the inconsistent and weak nature of the effects observed.

### 4.1. Naturalistic stimulus viewing elicits weak adaptation effects

Results of the current study suggest that adaptation to information embedded in naturalistic stimuli is relatively weak and inconsistent between adaptor orientations, as well as over the course of a single testing session. There are several elements of the current study that differ from typical adaptation investigations, which might contribute to the pattern of results found. First, the test stimulus in our experiment was relatively high contrast, which might act to limit adaptation effects (Parker, 1972). However, tilt aftereffects have been demonstrated using high-contrast stimuli (Gurbuz & Boyaci, 2023; Knapen et al., 2010; Nakashima & Sugita, 2014; Wolfe, 1984). Furthermore, the reader can experience the effect themselves in simple high contrast demonstrations, like that shown in **Fig 1A**. Hence, it is unlikely that such contrast-dependent adaptation could explain the current results.

Further, previous research has demonstrated clear adaptation effects within limited experimental sessions with as little as two minutes of adaptation (Magnussen & Johnsen, 1986). However, unlike past experiments employing minimalistic stimuli, the current task inherently allows visual input to change substantially across adaptation due to the inclusion of eye movements in conjunction with a dynamic film stimulus. It is possible that the use of a dynamic stimulus, as in the current study, inherently weakens adaptation effects and/or requires longer exposure times to elicit to the same extent as in past research.

### 4.2. The impact of image phase

While the current study manipulated the oriented contrast distributions across film frames to have a single orientation at relatively low spatial frequencies, and equated orientation energy at all other spatial frequencies. However, we did not alter the phase information within each frame. Phase alignment, therefore, will have preserved the spatial structure of the film frames (Hansen & Hess, 2007; Rideaux et al., 2022; Thomson & Foster, 1997; Zavitz & Baker, 2014). In the current study, stimulus processing still subjectively results in very strong orientation cueing from structural information such as the edges of buildings, regardless of the manipulated oriented contrast energy. Indeed, previous investigations have successfully had participants match naturalistic images based on their spatial structures after undergoing manipulation of the overall phase alignment and being filtered to have uniform oriented contrast (Hansen & Hess, 2007). As such, the preservation of spatial structure in the current study’s stimuli is expected to have allowed meaningful engagement with the film, facilitating following of the storyline and appreciation of the scenes they were observing. It is therefore possible that such scene “understanding”, particularly in the current study where global structural information is in alignment with naturalistic image statistics, might have mitigated adaptation effects. This possibility is an exciting question for future research.

### 4.3. The impact of participant’ stimulus viewing patterns

Given that participants freely viewed the video stimulus, it is likely that the peripheral visual field was inconsistently exposed to the adaptor stimulus, which is potentially relevant given the localised nature of adaptation effects (Gibson, 1937). Depending on a participant’s fixation distribution, parts of the visual field could be exposed to the adaptor stimulus while others would not. Further, it is possible that the areas that are exposed to the adaptor could align with the subsequent standard stimulus. This could be problematic due to the “El Grecco” effect, whereby a viewer who experiences a perceptual distortion for a given stimulus should experience that same distortion for their reproduction of that stimulus (Firestone & Scholl, 2016). As such, their reproduction should reflect the “objective” nature of the stimulus, and not reflect the experienced distortion. In the current study, if the adapted area of the visual field was exposed to both the standard and test stimulus, both would be equally impacted by adaptation, preventing measurement of any adaptation effects that are present. However, such an issue would necessitate that participants consistently foveate particular peripheral areas of the visual stimulus during adaptation. Previous research has demonstrated viewers have a “central” bias, whereby they tend to fixate the centre of a display, even when viewing live-action film stimuli (Dorr et al., 2010). Additionally, visual exploration of naturalistic images is largely driven by salient low-level features that can change drastically in spatial location in dynamic film stimuli (Carmi & Itti, 2006; Elazary & Itti, 2008; Itti, 2005; Zangrossi et al., 2021). Further, if participants were to fixate particular peripheral areas of the film on every trial, we should not have observed any significant adaptation results. Hence, it is unlikely that such concerns could entirely explain the adaptation effects observed.

While it is unlikely that concerns regarding how participants engage with the visual stimulus can explain the observed results, future research could improve upon the current study by directly addressing such concerns. This could be done by having participants maintain fixation for the duration of the experimental session – however, this would reduce the naturalistic viewing conditions imposed in the current study and would likely require eye tracking to enforce. If eye tracking were implemented, participants could watch the film stimulus in a gaze-contingent manner, which would ensure that the visual field being adapted is controlled for and the standard stimulus could be positioned to account for this. Alternatively, experiments employing virtual/augmented reality (VR/AR) could be beneficial to explore these phenomena. However, the benefit of VR/AR would be to fully immerse participants in an environment with altered image statistics. This would likely be problematic in the case of measuring adaptation because it would be difficult to design an adaptation task with a standard that was not equally impacted by the adaptor.

### 4.4. Adaptation to naturalistic stimuli is inconsistent

Overall, the current study suggests that we exhibit weak and inconsistent adaptation in response to orientation statistics conveyed by live-action film stimuli. Further investigation into the contribution of participants eye movements would elucidate the contribution of participants’ overt attentional deployment to the visual stimulus to observed adaptation results. Further, the current study may point to longer exposure times being required to elicit strong adaptation effects in the context of more naturalistic viewing conditions. Further investigations reflecting the current results, however, would suggest evidence of strong adaptation effects in the literature to date might be a product of the abstract/simple stimuli that have been used.

## Data availability

Behavioural data and GLMM code are available on the Open Science Framework: https://osf.io/5evns/?view_only=ba38508be394490e85804691cd2f67e2

## Acknowledgements

We would like to thank Laura Wang for her assistance with data collection. This work was supported by Australian Research Council Discovery Early Career Researcher Awards awarded to WJH (DE190100136) and RR (DE210100790), as well as a National Health and Medical Research Council (Australia) Investigator Grant awarded to JBM (GNT2010141). RR was also supported by a National Health and Medical Research Council (NHMRC; Australia) Investigator Grant (2026318). EJA was supported by the Deutsche Forschungsgemeinschaft (DFG, German Research Foundation) – SFB/TRR 135 (project no. 222641018, project C1).

## Additional Information

The authors have no conflicts of interest to declare.

