## Supplemental Materials for "Investigating orientation adaptation following naturalistic film viewing"

Emily J A-Izzeddin[1,2*], Reuben Rideaux[2,3], Jason B Mattingley[2,4], William J Harrison[2,4,5]

[1] Department of Experimental Psychology, Justus Liebig University Giessen, Giessen, Germany

[2] Queensland Brain Institute, The University of Queensland, St Lucia, Queensland 4072 Australia

[3] School of Psychology, The University of Sydney, Camperdown, New South Wales 2006 Australia

[4] School of Psychology, The University of Queensland, St Lucia, Queensland 4072 Australia

[5] School of Health, University of the Sunshine Coast, Sippy Downs, Queensland 4556 Australia

### S.1. Overview of experimental sessions

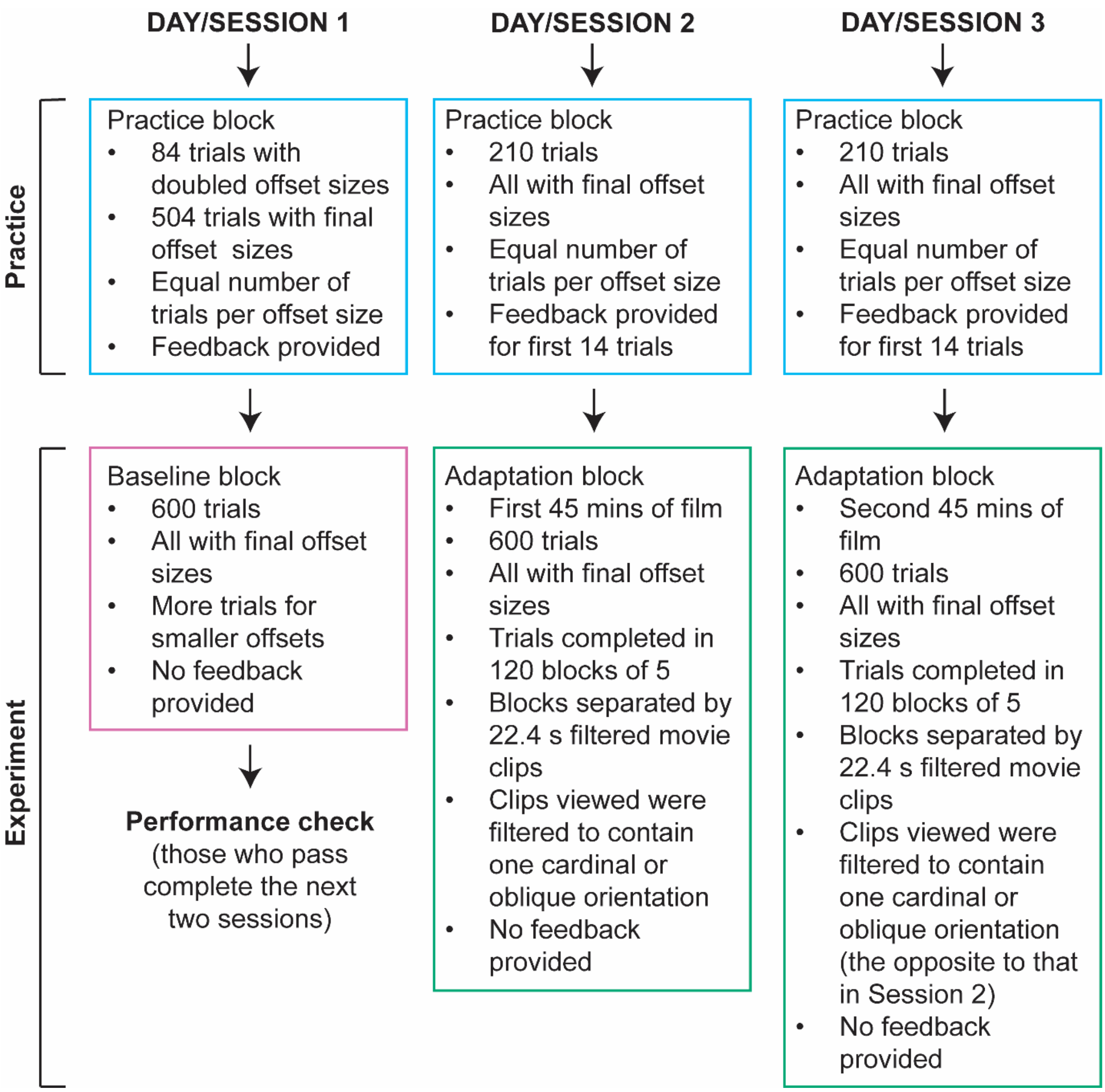


**Figure S.1. Overview of experimental sessions.** Each participant completed an initial session (left column) that included a practice block of trials (blue box), followed by the baseline measurement (pink box). Participants' baseline performance is assessed (see Practice and baseline testing in the main text). Participants who pass the baseline performance assessment go on to complete the next two sessions (middle and right column). Each of the final two sessions followed the same structure: participants initially did a block of practice trials (blue boxes), followed by an adaptation block (green boxes). In the adaptation blocks, participants performed the same behavioural task as in the baseline session. However, in the adaptation blocks, trials were completed in blocks of five, separated by a filtered movie clip, containing an adaptor that was either cardinal (0° or 90°) or oblique (45° or 135°). Each participant did one session with a cardinal adaptor and the other with an oblique adaptor. The order of cardinal vs oblique adaptors was counterbalanced across participants, as were combinations of the possible adaptor orientations.

### S.2. Full GLMM output

**Table S.1: Full output for the GLMM defined by the equation:** $\boldsymbol{y \sim \beta_0 + \beta_1 T + \beta_2 F + \beta_3 S + \beta_4 FS}$**.** Here, $\boldsymbol{\beta_0}$ is the intercept term, $\boldsymbol{\beta_1}$ is the weight of the test stimulus offset relative to the standard, $\boldsymbol{T}$, $\boldsymbol{\beta_2}$ is the weight of the adaptor condition, $\boldsymbol{F}$, $\boldsymbol{\beta_3}$ is the weight of the standard orientation, $\boldsymbol{S}$, and $\boldsymbol{\beta_4}$ is the weight of the adaptor/standard orientation interaction, $\boldsymbol{FS}$. To partially pool coefficient estimates across participants, the GLMM included participant as a random effect.

| *Name* | *Estimate* | *SE* | *tStat* | *DF* | *pValue* |
|---|---|---|---|---|---|
| *Intercept* | 0.065 | 0.090 | 0.726 | 52587 | 0.468 |
| *F_0* | -0.067 | 0.095 | -0.708 | 52587 | 0.479 |
| *F_45* | 0.050 | 0.102 | 0.497 | 52587 | 0.620 |
| *F_90* | -0.104 | 0.145 | -0.718 | 52587 | 0.473 |
| *F_135* | 0.037 | 0.128 | 0.292 | 52587 | 0.771 |
| *S_-22.5* | -0.414 | 0.136 | -3.048 | 52587 | 0.002* |
| *S_22.5* | 0.090 | 0.170 | 0.531 | 52587 | 0.596 |
| *S_67.5* | -0.115 | 0.129 | -0.892 | 52587 | 0.373 |
| *T* | -0.275 | 0.018 | -15.081 | 52587 | <.001** |
| *F_0:S_-22.5* | 0.567 | 0.130 | 4.350 | 52587 | <.001** |
| *F_45:S_-22.5* | -0.191 | 0.141 | -1.355 | 52587 | 0.176 |
| *F_90:S_-22.5* | -0.004 | 0.168 | -0.022 | 52587 | 0.983 |
| *F_135:S_-22.5* | 0.099 | 0.164 | 0.604 | 52587 | 0.546 |
| *F_0:S_22.5* | -0.033 | 0.170 | -0.194 | 52587 | 0.846 |
| *F_45:S_22.5* | 0.190 | 0.207 | 0.921 | 52587 | 0.357 |
| *F_90:S_22.5* | 0.026 | 0.192 | 0.134 | 52587 | 0.894 |
| *F_135:S_22.5* | 0.064 | 0.131 | 0.491 | 52587 | 0.623 |
| *F_0:S_67.5* | -0.050 | 0.182 | -0.277 | 52587 | 0.782 |
| *F_45:S_67.5* | -0.232 | 0.193 | -1.202 | 52587 | 0.229 |
| *F_90:S_67.5* | 0.336 | 0.165 | 2.036 | 52587 | 0.042 |
| *F_135:S_67.5* | -0.389 | 0.234 | -1.665 | 52587 | 0.096 |

* $p < .005$

** $p < .001$
